# MicrobeTrace: Retooling Molecular Epidemiology for Rapid Public Health Response

**DOI:** 10.1101/2020.07.22.216275

**Authors:** Ellsworth M. Campbell, Anthony Boyles, Anupama Shankar, Jay Kim, Sergey Knyazev, William M. Switzer

**Affiliations:** Centers for Disease Control and Prevention, Atlanta, GA 30329; Northrup Grumman, Atlanta, GA 30345; Oak Ridge Institute for Science and Education, Oak Ridge, TN 37830; Department of Computer Science, Georgia State University, Atlanta, GA, 30303

## Abstract

**Motivation:** Outbreak investigations use data from interviews, healthcare providers, laboratories and surveillance systems. However, integrated use of data from multiple sources requires a patchwork of software that present challenges in usability, interoperability, confidentiality, and cost. Rapid integration, visualization and analysis of data from multiple sources can guide effective public health interventions.

**Results:** We developed MicrobeTrace to facilitate rapid public health responses by overcoming barriers to data integration and exploration in molecular epidemiology. Using publicly available HIV sequences and other data, we demonstrate the analysis of viral genetic distance networks and introduce a novel approach to minimum spanning trees that simplifies results. We also illustrate the potential utility of MicrobeTrace in support of contact tracing by analyzing and displaying data from an outbreak of SARS-CoV-2 in South Korea in early 2020.

**Availability and Implementation:** MicrobeTrace is a web-based, client-side, JavaScript application (https://microbetrace.cdc.gov) that runs in Chromium-based browsers and remains fully-operational without an internet connection. MicrobeTrace is developed and actively maintained by the Centers for Disease Control and Prevention. The source code is available at https://github.com/cdcgov/microbetrace.

## 1. Introduction

The burgeoning field of public health bioinformatics has given rise to a plethora of specialized software for analysis and visualization of pathogen genomic data to aid outbreak investigations (Clément, et al., 2018; Leipzig, 2017). Implementation of these analytic tools can be complex and fraught with a variety of technical and administrative barriers, like faulty install procedures or the need for administrative credentials to install (Sussman, 2007). As a result, routine use of bioinformatic tools in public health can be delayed or blocked because users lack the wide range of skills necessary to install, operate, and integrate them (Pond, et al., 2018). Historically, many public health workers with educational backgrounds in medicine, epidemiology, and laboratory sciences lack informatics skills needed to collect, analyze and display data (*Applications of Clinical Microbial Next-Generation Sequencing: Report on an American Academy of Microbiology Colloquium held in Washington, DC, in April 2015*, 2015). This skill mismatch tends to be more pronounced at local health departments, representing the frontlines of public health, which have limited capacity and funding for informatics, cyber security, and computational infrastructure (Gwinn, et al., 2017).

The complex landscape of public health bioinformatics has necessitated the development of tools designed to sidestep hurdles that can hinder adoption or routine use. Technical and administrative barriers are often reduced by moving complex analytics and computation to off-site servers. However, while cloud computing has revolutionized the healthcare industry (Celesti, et al., 2019), state public health laws often prohibit the storage of sensitive data on off-site servers in the cloud. Tool accessibility can also be hampered by cluttered user interfaces (Bastian, et al., 2009; Hall, 1999; Maths, 2007; Smoot, et al., 2011) and unwieldy workflows that hamper human-computer interaction (Argimón, et al., 2016; Hadfield, et al., 2019; Hadfield, et al., 2018; Pond, et al., 2018). Given the breadth of genetic sequencing technologies and bioinformatic methods, tool adoption can suffer when acceptable input and output file formats are limited, complicating or even preventing integration with existing systems and workflows. To foster adoption and routine use, bioinformatic tools should be secure, easy to use, and capable of accepting or exporting data in commonly used formats.

To accommodate the specific needs of local health departments, we developed a standalone but browser-based tool to integrate, visualize and explore data routinely collected during public health investigations of outbreaks and transmission clusters. These data can include case lists describing demographic and behavioral information, case lists with high-risk contacts, in addition to pathogen genomic data. MicrobeTrace was designed to enable users to construct pathogen genetic distance networks and visually integrate them with contact tracing networks to better characterize a transmission network. MicrobeTrace users can further characterize their integrated networks by mapping additional metadata to visual attributes like size, shape and color. In contrast with other tools commonly used for transmission analysis (Argimón, et al., 2016; Hadfield, et al., 2018), all visual attributes can be modified by the user via simple interactions (e.g., dropdown menus, toggle buttons, and color pickers) in real-time, without modification of the underlying data. MicrobeTrace is well suited for working with personally identifiable information (PII) because it performs all computations and visualizations on the user’s computer and does not store or transmit any data from the user’s computer. When using a supported and updated web browser (e.g., Chrome, Firefox, or Edge) all cached files are cleared when the browser session ends unless caching is explicitly enabled by the user. At no time are user data transmitted anywhere over the internet. As a result, MicrobeTrace can be accessed from the CDC website initially and thereafter used with data stored on the user’s computer without an internet connection, making it ideal for rapid visualization of data in the field.

Here, we present MicrobeTrace and describe its utility across multiple public health use cases including retrospective analyses and outbreak response. We also report on its use in transmission analysis for a broad spectrum of infectious diseases, such as tuberculosis, viral hepatitis, sexually transmitted diseases as well as special pathogens like SARS-CoV-2 and Ebola.

## 2. Methods

### 2.1 Development

MicrobeTrace has been developed according to an agile and open source model, with all code available via GitHub.com (Boyles and Kim, 2018). This enables users to directly observe the rate of development as well as submit and monitor feature requests and system bug reports. MicrobeTrace development has been guided by requirements and features requested by public health practitioners who will use the application in their routine field work. All code is indexed by the federal open source repository (Code.gov, 2019) and promoted by Code.gov (Code.gov, 2019). The MicrobeTrace codebase is regularly scanned by Fortify Software (HP Enterprise Security Products, 2020) and SonarQube (SonarQube.org, 2020) to ensure security and code stability. Further, all related modules of code that depend on each other are automatically monitored for vulnerabilities and updated by GitHub’s Dependabot service. This automated monitoring service ensures that security vulnerabilities are rapidly detected, reported to our development team, and addressed. GitHub’s *Actions* service is used to automate the process of testing newly developed features before official release. This process of automated testing ensures that each time new features are added into MicrobeTrace, all pre-existing functionality are automatically tested prior to an official release.

### 2.1 Outreach

Training and outreach are important factors in refining a software product through interaction with the user base. Training is provided through three modalities: (1) small ad-hoc webinar sessions (5-20 attendees) to support specific outbreak and cluster investigations, (2) large in-person training sessions (20-100+), and (3) a recorded webinar available via YouTube (CDC, 2020) that is compliant with Section 508 of the Rehabilitation Act of 1973. A detailed, 508-compliant >100-page manual is also available for download on the GitHub website (Shankar, et al., 2019). Finally, a brief ‘flyer’ describing the tool’s general functionality (Campbell, 2019) is available in PDF format, for handout at public health and academic conferences.

## 3. Results

### 3.1 Data Formats

MicrobeTrace handles a variety of file types and formats that are traditionally collected during public health investigations. Pathogen genomic information can be integrated as raw genomic sequences, genetic distance matrices, pairwise genetic distances, or phylogenetic trees. Epidemiologic and other metadata about cases (node lists) and their high-risk contacts (edge or link lists) can be integrated as spreadsheets. Importable in a variety of file formats, these file types can be visualized independently or in-concert to achieve different analytic goals (Fig. 1). Early in an outbreak investigation, high-risk contacts can be combined with other epidemiologic information to visualize and characterize a risk network. When genomic data become available later in the investigation, genetic networks can be integrated to visualize concordance between epidemiologic and laboratory data sources. Alternatively, all available data sources can be integrated to construct a more holistic visualization of an ongoing public health investigation.

**Figure 1:**
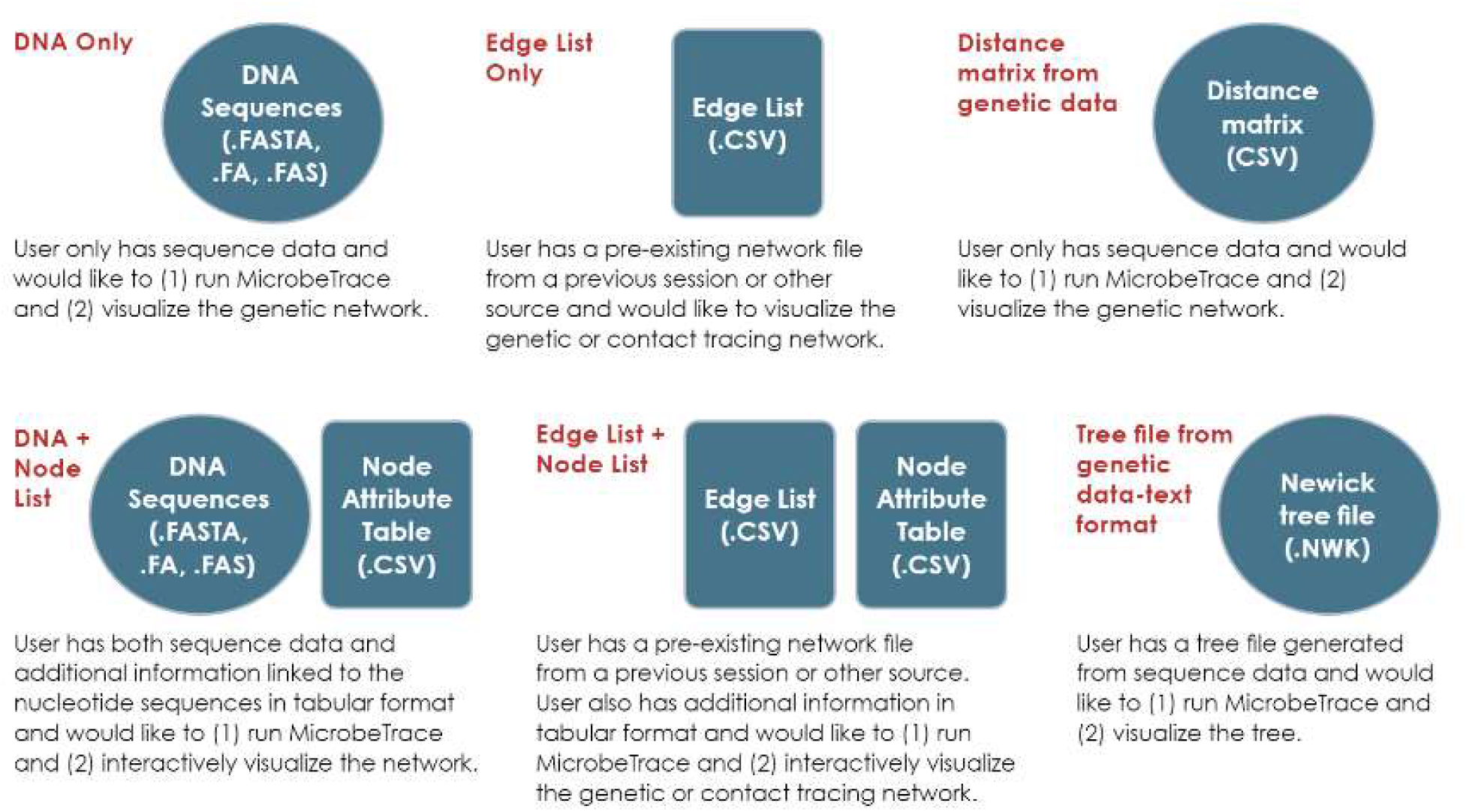
MicrobeTrace accepts input data in a variety of formats. This figure displays the most common use cases and their required files.

### 3.2 Preserving Data Security and Confidentiality

The information processing technology within MicrobeTrace is well adapted for use in a public health setting because it prioritizes the confidential but effective use of sensitive data collected during an outbreak investigation. MicrobeTrace was developed as a *client-side only* application that is incapable of transmitting any user data over the internet. In contrast, most web-based bioinformatic applications require the user’s data be submitted over the internet for processing by a remote *server-side* application before results can be returned to the user. Local processing is achieved through open source development and translations of traditional bioinformatic algorithms to align (Boyles, 2019a; Li, 2014; Smith, et al., 1981), compare (Boyles, 2019b; Pond, et al., 2018; Tamura and Nei, 1993), and evaluate genomic sequences and their relationships to one another (Boyles, 2019d; Fourment and Gibbs, 2006; Knyazev, 2020; Kruskal, 1956). Importantly, sequence (a) alignment, (b) comparisons, (c) phylogeny, and (d) network evaluations are recapitulations of established methods and do not constitute novel development. Therefore, to the best of our knowledge, the results derived from these JavaScript methods are interchangeable with results derived from their respective, native implementations. A novel extension of the network evaluation method is described below in section 3.4 as the ‘Nearest Connected Neighbor’.

Visualizations must be generated with care during an outbreak investigation to ensure confidential and narrow use of sensitive data. PII and other sensitive information like geospatial coordinates, zip codes, and phone numbers should only be accessible to Disease Investigation Specialists conducting contact tracing interviews. However, an epidemiologist performing a retrospective analysis can use the same visualization layout with remapped labels, colors, shapes and sizes. Indeed, sensitive geocoordinates can still be used confidentially to produce informative maps by applying the random ‘jitter’ function in MicrobeTrace to reduce the precision of the displayed map marker. In concert, these diverse and accessible controls enable public health experts to safely and confidently leverage sensitive data without risk to the public’s confidentiality.

### 3.3 Genetic Distance Networks

To demonstrate the bioinformatics capacity of MicrobeTrace, we used a publicly available HIV-1 data set consisting of 1,164 sequences of the partial polymerase (*pol*) region (GenBank accession numbers KX465238-KX467180) from a recent study in Germany in addition to associated metadata describing behavioral risk factors and gender (Pouran Yousef, et al., 2016). Partial pol sequences are typically collected for determination of antiretroviral drug resistance monitoring for care and treatment for persons living with HIV infection.

The bioinformatics workflow of genetic distance networks in MicrobeTrace begins with a pairwise sequence alignment of each input sequence against a reference, according to the Smith-Waterman algorithm (Boyles, 2019a; Li, 2014; Smith, et al., 1981). Multiple sequence alignments are too time constrained and are not used. A user can align to a curated reference, an arbitrary custom reference, or the first input sequence. For HIV-1, the strain HXB2 from the United States (U.S.) is a common reference sequence (GenBank accession number K03455). Once aligned, pairwise genetic distances are calculated according to either a raw hamming distance or the Tamura-Nei substitution model (TN93) (Boyles, 2019d; Pond, et al., 2018; Tamura and Nei, 1993). When the TN93 substitution model is selected, handling of ambiguous bases can be configured as previously described (Pond, et al., 2018). Pairwise genetic distances can be easily filtered by a threshold defined by the user, in this case 1.5% nucleotide substitutions per site (Fig 2A). Notably, users are empowered with the tools necessary to identify and select the distance threshold value that best fits their public health use case (Wertheim, et al., 2017). In some situations for HIV-1, a conservative threshold of 1.5% genetic distance might be appropriate to best understand the historical evolution of recent transmission events (Wertheim, et al., 2014). A more stringent TN93 threshold of 0.5% is often used to identify the most recent and rapid clusters of HIV-1 transmission (Fig 2B). Threshold determinations are often informed by cluster size and growth rate criteria (Erly, et al., 2020; France and Oster, 2020; Oster, et al., 2018). MicrobeTrace offers the ability to filter by genetic distance and cluster size thresholds in the same ‘Global Settings’ menu. Here, using the German HIV-1 dataset we have filtered for clusters of size N ≥ 5 after the 1.5% genetic distance threshold is applied. This filter hides 73.1% (N = 851) of individuals that are too genetically distant to cluster with any other sequences in the sample as well as 17.9%(N = 208) of individuals whose HIV-1 sequences reside in clusters of size N ≤ 4. HIV-1 sequences from the remaining 9.0% (N = 105) of individuals are displayed as genetic distance networks in Figure 2. Variables of interest can be readily mapped to the nodes or links, including HIV-1 pol drug resistance mutations to identify clusters of transmitted drug resistance (Fig 2C).

**Figure 2:**
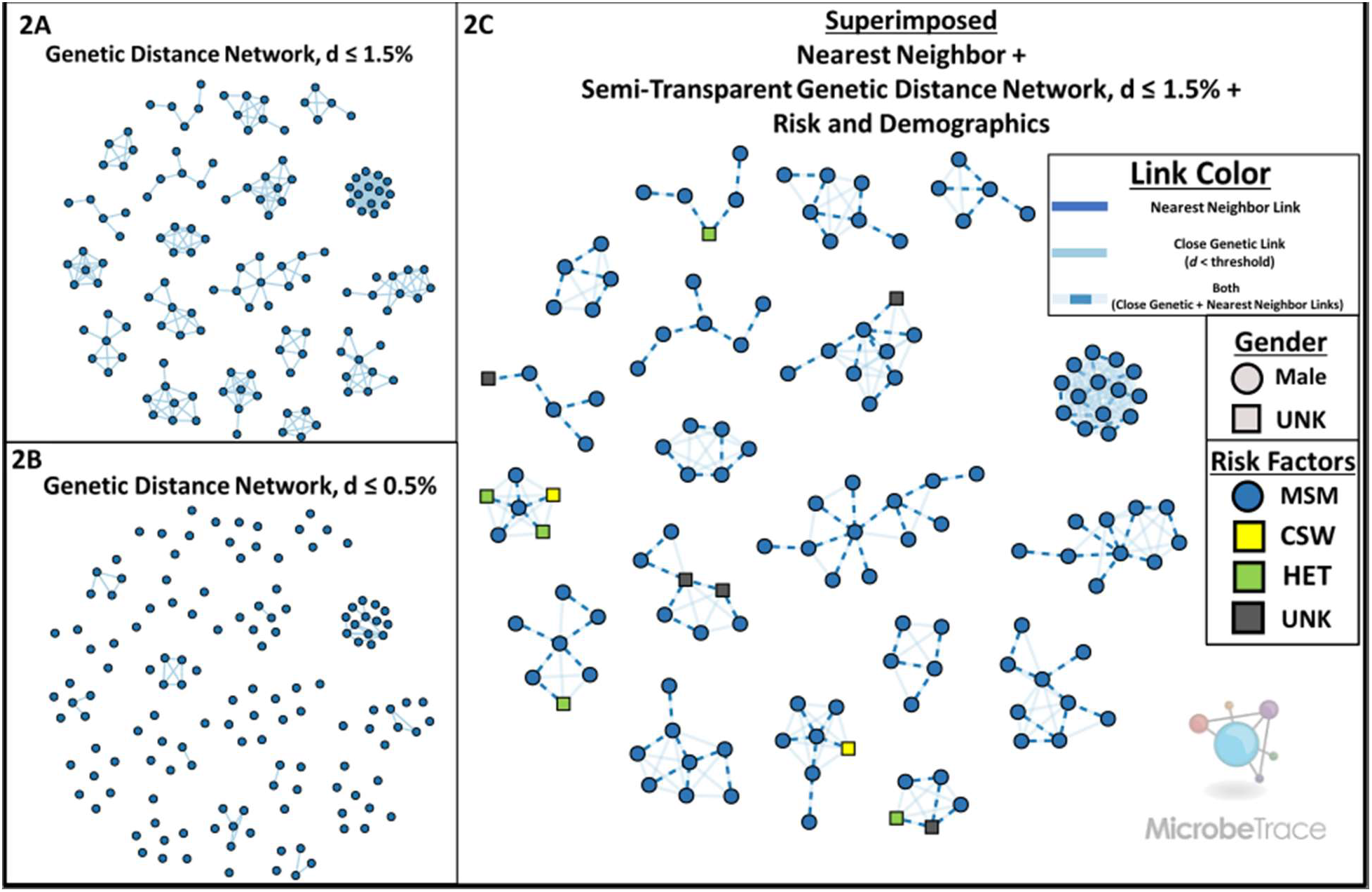
MicrobeTrace excels at rendering pathogen genetic distance networks and mapping visual characteristics to user-provided metadata. (**2A**) The HIV-1 partial polymerase (pol) distance network, with a genetic distance threshold (d) of 1.5%. (**2B**) The same HIV-1 pol network shown in 2A with node positions held constant, but with a more stringent genetic distance threshold (d) of 0.5%. (**2C**) The same HIV-1 pol network shown in 2A with node positions held constant. Nearest connected neighbor links have been superimposed as dashed lines. The transparency of links that do not connect nearest neighbors has been increased. Gender and transmission risk factors have been mapped to node shape and color, respectively.

### 3.4 Arbitrary Genetic Distance Networks

A simple nucleotide substitution model is not always suitable to understand phylogenetic relationships. Rather than require the use of a single model, MicrobeTrace supports the integration of precomputed distance matrices and pairwise distance lists. A user can provide any pre-computed pairwise distances, regardless of the underlying nucleotide substitution model, as a list or a matrix in order to render those data as a network. For distance matrices, both full matrix and PHYLIP formats are accepted. MicrobeTrace also provides a novel and simple filtering algorithm to render only the nearest connected genetic neighbor(s) for each node, while still maintaining cluster connectivity. Where any two genetically equidistant neighbors are possible, both links are rendered when the ‘Nearest Connected Neighbor’ filter is applied. This approach is particularly useful to understand the historical context of an entire cluster, while focusing on the part of the cluster exhibiting the most concerning and rapid growth. For example, an HIV cluster in rural southeastern Indiana grew rapidly in 2015 but underwent slow growth for nearly a decade prior (Campbell, et al., 2017). The nearest connect neighbor method yields results similar to a non-exhaustive search for all minimum spanning trees, as has been previously described (Bbosa, et al., 2020; Campbell, et al., 2017). The threshold and nearest connected neighbor filters are not mutually exclusive and can therefore be applied simultaneously to ensure that genetically distant nodes remain disconnected. This enables the inclusion of related, but more distant sequences in a cluster visualization while minimizing the information overload typically accompanied by increased distance thresholds (as shown in Fig. 2A). HIV-1 genetic distance links that fell below the 1.5% threshold but were not included as a nearest connected neighbor link are shown at reduced opacity (Fig. 2C).

### 3.5 Patristic Distance Networks

Phylogenies are ubiquitous in public health and bioinformatics, but a phylogeny may be difficult to integrate with more traditional contact tracing data. While powerful new tools are available to integrate taxa-level characteristics into phylogenies, integration of paired contacts is unavailable. Instead, the genetic distances encoded on the phylogeny must be measured and recast as pairwise patristic distances of a phylogeny. Specifically, these are tip-to-tip measurements between individuals on an evolutionary tree that account for the most recent common ancestor. This step is necessary, because it results in a pairwise genetic distance list that is readily integrated with pairwise contact data. Provided a phylogenetic tree in Newick format, MicrobeTrace will traverse the phylogeny to calculate and render the pairwise patristic distance network corresponding to that phylogeny.

### 3.6 Epidemiologic Networks

Importantly, phylogenies or pathogen genetic sequence data are not required to leverage MicrobeTrace to visualize public health data. MicrobeTrace supports the visualization of arbitrary networks, such as those collected during contact tracing during an outbreak or cluster investigation. Acceptable networks are not limited to person-to-person links but can include person-to-place or place-to-place. To visually differentiate persons from places, MicrobeTrace can style the shape of any network node according to a node type column (e.g., nodeType = ‘Person’ or ‘Place’) defined in the data set. If additional metadata are available to describe a link, it can be colored according to user-defined categorical variables. Alternatively, an option is provided to scale link width according to a user-defined numeric variable or its reciprocal.

### 3.7 Multi-Layer Networks

Epidemiologic and genetic networks often offer complementary perspectives about transmission clusters (Campbell, et al., 2020). MicrobeTrace can render an arbitrary number of networks simultaneously by representing multiple overlapping links between pairs of nodes (e.g., hyperlinks) as color-mapped, dashed lines. In addition to independent color-mappings according to underlying data, the effect of a particular network layer can either be hidden or accentuated via independent transparency controls. For example, to protect individual privacy, public health experts may choose to make epidemiologic reports of high-risk contact invisible while rendering only close genetic links when producing figures for public consumption.

### 3.8 Maps with Network Overlay

Integrated epidemiologic and genetic networks are abstract diagrams that can be used to inform policy and prevention efforts when augmented with additional information. MicrobeTrace can generate choropleth maps, globe diagrams, or more common map projections. MicrobeTrace mapping functions offline with pre-computed shapefiles describing countries, as well as U.S. states and counties. Should internet access be available, MicrobeTrace can be configured to request high-resolution geospatial map tiles from a JavaScript map service called Leaflet (Agafonkin, 2014). MicrobeTrace also enables users to contextualize their maps with a network overlay that maintains all color mappings defined in the network visualization. Users can select from various geographic units, ranging from Country, and – at present – state, county, and zip codes for the U.S. or paired latitude and longitude values. For each geographic level, a marker is placed at the geographic centroid. Over-plotting can be addressed by a combination of automated aggregation or manual transparency tools. Maps can also be customized with user-provided geospatial data in the GeoJSON format.

### 3.9 Customization and Interactive Exploration

To demonstrate the generalized visualization capacity of MicrobeTrace, we present a publicly available data set describing clinical, demographic and contact tracing data derived from the Korean Centers for Disease Control (KCDC) investigation of the COVID-19 outbreak (Kim, 2020). The data set does not contain coronavirus sequence data, but instead details 383 transmission histories between 510 cases. It also contains an additional 1,627 cases of COVID-19 with no documented transmission histories. As before, using filtering capabilities unique to MicrobeTrace, we limit our visualizations to transmission clusters of size ≥ 5 cases (Fig. 3).

**Figure 3:**
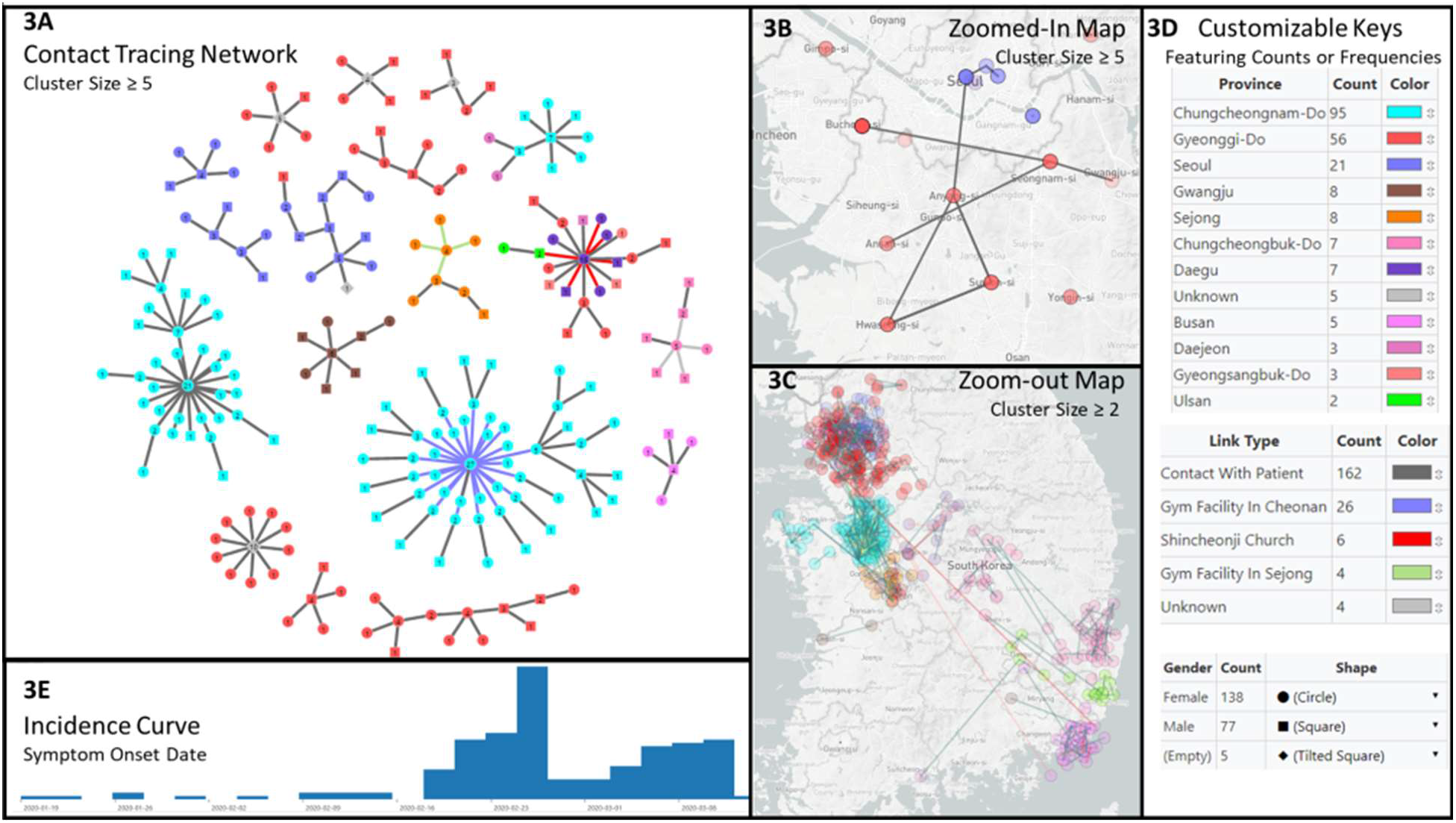
MicrobeTrace allows the creation of informative dashboard visualizations. (**3A**) Reports of high-risk contact between COVID-19 cases in clusters of size N ≥ 5, nodes are (i) colored by province, (ii) shaped by gender, and (iii) labeled with the total number of high-risk contacts. (**3B**) Geospatial map of clusters of size N ≥ 5 zoomed to show only Seoul, South Korea. (**3C**) Geospatial map of clusters of size N ≥ 2. Node positions have been randomly altered, via MicrobeTrace’s ‘jitter’ functionality, to preserve patient privacy. (**3D**) In-application color and shape keys that offer interactive color-pickers and labeling. (**3E**) Incidence curve showing symptom onset date.

MicrobeTrace is centered around integration and visualization of pathogen genomic and network data but is accompanied by an array of customizable tables, charts, and geospatial maps that facilitate exploration and communication of public health data. Each view is interactive and interoperable so nodes in one view are propagated to other tiled views. For example, a node selected by search or click in the **Table View** is highlighted both there and in relevant adjacent views. Similarly, all choices on color-mappings for nodes and links are propagated to all relevant adjacent views. All views are resizable and can be tiled to produce rich, interactive and exploratory dashboards as demonstrated below. We have tiled the COVID-19 transmission network, the symptom onset incidence curve, and a geospatial map with transmission network overlay (Fig. 3). Here, we perform the following visual manipulations within MicrobeTrace: (1) automatically calculate and map the number of contacts for each case to the label that is centered over each node (Fig. 3A), (2) map the node color to the case’s province (Fig. 3A-D), (3) map link color to the mode of exposure (Fig. 3A-D), (4) map node shapes to the case’s gender (Fig. 3A) (5) superimpose the network onto a high-resolution geospatial 2D map projection (Fig. 3B-C), (6) tailored color, size and transparency to desired values (Fig. 3B-C), and (7) generated an incidence curve according to the date of symptom onset (Fig 3E).

As with genetic data, networks are not required to leverage most of the visualizations in MicrobeTrace. Indeed, MicrobeTrace can be used to achieve rich visualizations using a list of nodes with a handful of variables like *age, gender, province, city, exposure type, symptom onset date, test confirmation date* and *hospital release data*. We demonstrate the construction of complex figures like a **Flow Diagram, Gantt Chart, Cross-tabulation, Aggregation, and Histogram** with simple dropdown menus (Fig. 4). Additional diagrams can be achieved with the **2D Network, 3D Network, Scatter Plot, Heatmap, Bubbles, Choropleth,** and **Globe Views** with relevant data types selected with simple dropdown menus. Operation of each view is documented in detail in the MicrobeTrace user manual (Shankar, Campbell, *et al*., 2019).

**Figure 4:**
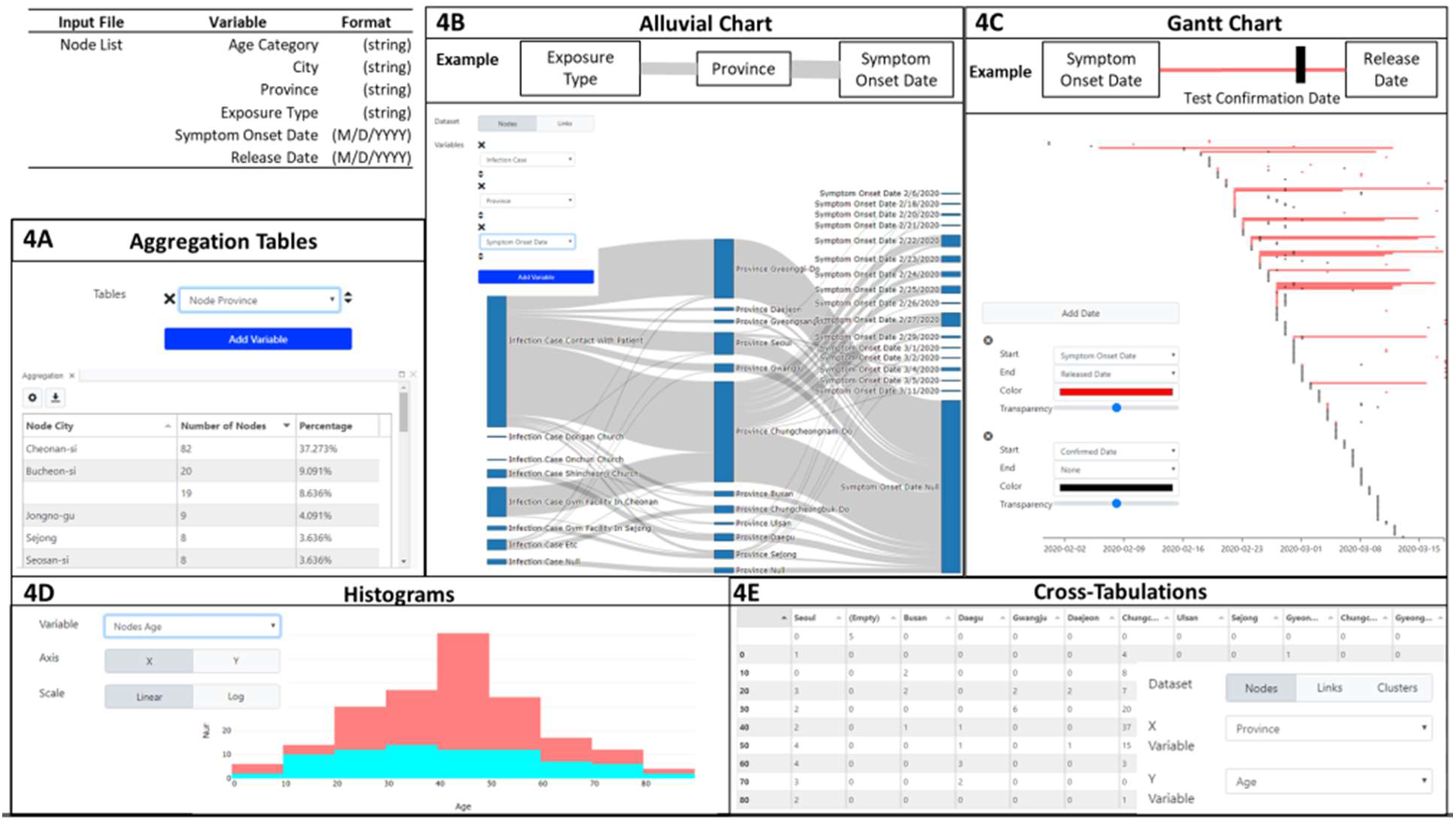
MicrobeTrace visualization does not require genomic or contact tracing data and calculate aggregation and cross-tabulation tables in addition to visualizing histograms, alluvial/flow diagrams and Gantt charts. Each diagram has an inset settings menu that describes the settings changes necessary to achieve them. (**4A**) City-level aggregation achieved via a single dropdown selection. (**4B**) Alluvial diagram of associations between the Type of Exposure to COVID-19, Province, and Symptom Onset Date. (**4C**) Gantt charts to describe the span of time between Symptom Onset, Positive Test Confirmation, and Hospital Release Date. (**4D**) Age histogram, binned by decade and colored by gender. This histogram illustrates a trend identified during the early Korean outbreak, wherein a disproportionate number of middle-age female cases was diagnosed (**4E**) Cross-tabulation table of cases by City and Age categories.

### 3.10 Sequence alignment and phylogenetic tree views

When sequence data are available, a variety of additional diagrams and views are available. For example, the **Sequences View** can be used to export or check the quality of the pairwise alignment. The **Phylogenetic Tree View** will construct a tree via a neighbor-joining algorithm according to the provided pairwise distance calculations. The **Phylogenetic Tree View** has robust customization controls that have been modularized in a separate JavaScript library called TidyTree (Boyles, 2019c).

### 3.11 Reproducibility

Public health investigations are iterative and the underlying data sources tend to grow over time. Once MicrobeTrace workspaces have been customized they can be saved in two ways: (1) as a custom. MicrobeTrace file or (2) as a “stashed” (cached) browser session. As new data arrives, a user can choose to add new files and recompute the network while pinning nodes to their original positions on-screen. This capability enables a greater understanding of transmission dynamics by enforcing continuity between visualization and exploration sessions over time. Styling parameters and custom visualizations can be stored *independently* from the underlying data as a MicrobeTraceStyle file to facilitate communication between collaborators and preserve confidentiality. Style files can also be used to ensure continuity between public health investigations, such that different investigations yield identically styled visualizations even with different underlying data.

### 3.12 Data and visualization exports

Communicating data arising from public health investigations is a complex process that requires many fine adjustments, as messages are tuned to their audiences. To meet this need, MicrobeTrace is designed to provide users maximum control over visualization customization and export capabilities. For example, communication to academic and public health audiences often involves poster presentations that require images be scaled-up for large printer formats. We accommodate this requirement by enabling users to set specific export resolutions for PNG and JPEG formats. Alternatively, visualizations can be exported as Scalable Vector Graphics (SVGs) that can be enlarged to any arbitrary size without a loss of resolution. By default, a MicrobeTrace watermark is placed on images exported from MicrobeTrace; however, the transparency of the watermark can be increased using a menu slider to render it invisible. Taken together, these capabilities offer publication-ready image exports for scientific journals.

MicrobeTrace maximizes interoperability with other applications by enabling the export of all calculated and integrated datasets. The **Table View** renders tabular data which can be exported to comma-separated (CSV) and Excel (XLS, XLSX) formats. The node-level table includes all information joined from multiple input data sources as well as calculated fields like a node’s number of neighbors (‘degree’) and its cluster ID. The link-level table also includes calculated fields; for example, whether a link was identified as a ‘nearest connected neighbor’ as a Boolean result. MicrobeTrace offers robust filtering and selection capabilities that are also reflected in exported tables, ‘Selected’ and ‘Visible’ states are shown as Boolean results. Tables produced in the **Aggregation View** can be exported as formatted PDFs, CSVs, a zipped collection of CSVs, or an XLS/XLSX workbook where each aggregation is shown on independently named worksheets (Fig. 4A). Data derived from the **Map, Globe,** and **Choropleth Views** can be exported as GeoJSON files for interoperability with other Geographic Information System (GIS) software. Genomic sequence alignment can be exported in the FASTA or MEGA file formats in the Sequences View.

### 3.13 Statistics and analysis of MicrobeTrace usage

While some public health investigations that leveraged MicrobeTrace have been reported in the academic literature, many use cases supporting public health missions are never intended for publication or dissemination (Cranston, et al., 2019; Hogan, et al., 2017; John, et al., 2019; Shankar, et al., 2019; Falade-Nwulia, et al., 2018). To better understand that broad base of engagement, MicrobeTrace usage statistics are captured and reported by region via Google Analytics. When MicrobeTrace is accessed while the user is online, an anonymous Google Analytics cookie is sent along with information about the user’s rough geolocation and usage time. It is important to note that, offline usage is not tracked by Google Analytics. Since the launch of MicrobeTrace in March 2018, 2,642 unique users have connected for a total of 6,501 sessions (2.46 sessions per user) for a combined 738.6 hours of use (6.8 per session and 16.7 minutes per user). The overwhelming majority of users connect from the U.S. (N = 2,323, 87.8%) with the most prevalent international use coming from China (N = 55, 2.1%), the United Kingdom (N = 38, 1.4%), and Vietnam (N = 30, 1.1%). 50 additional countries account for the remaining 6.6% (N = 196) of users. Usage increases on weekdays, as the public health workforce goes to work, and the mean number of weekday users has increased from 1.1/weekday in February 2018 to highs of 20.5 and 14.6 per weekday in February and March 2020, respectively. (Fig. 5). Notably, as much of the world’s public health workforce has turned its attention to COVID-19 in February and March of 2020, MicrobeTrace usage peaked (Fig. 5).

**Figure 5:**
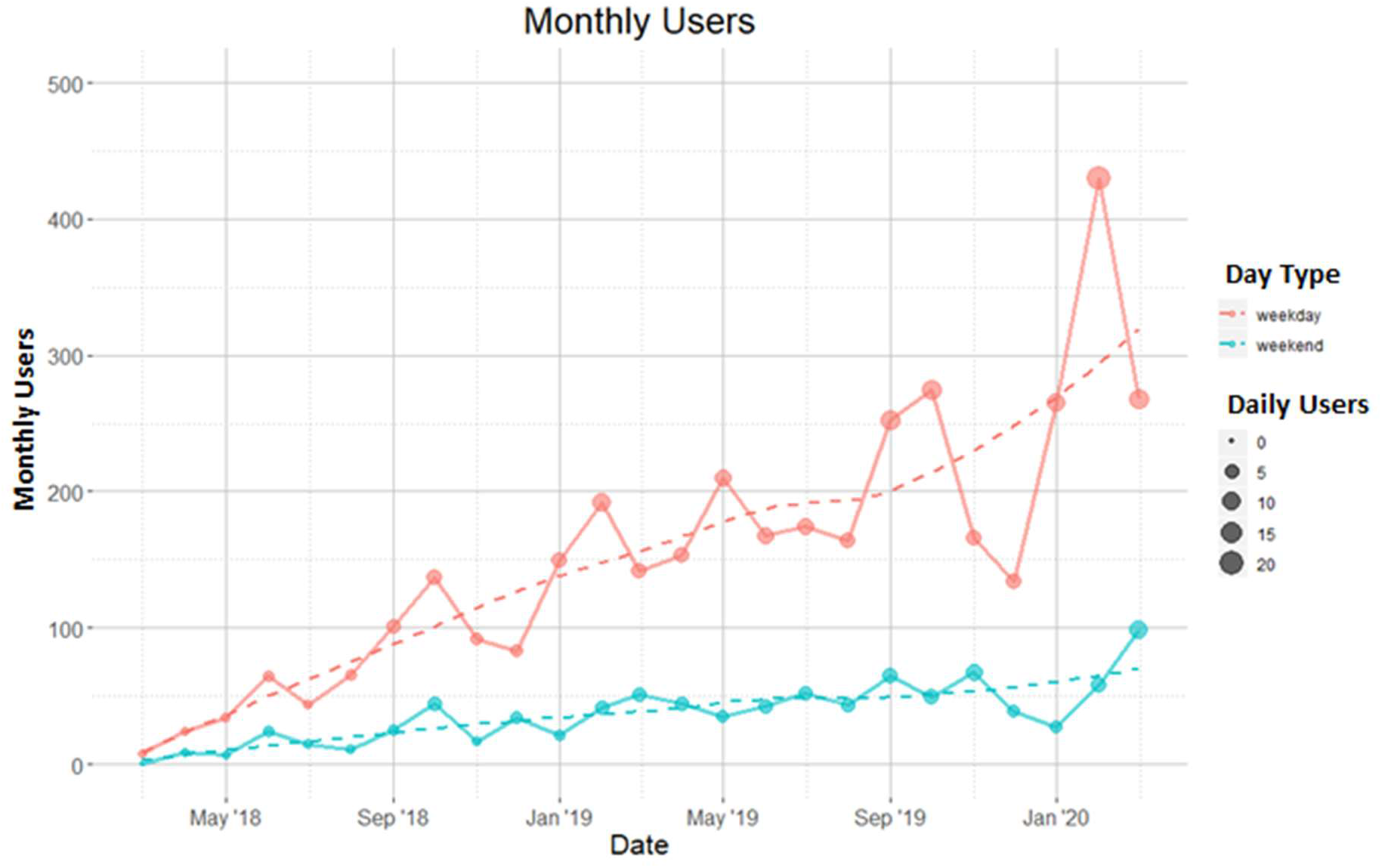
MicrobeTrace’s primary user base are public health officials during the work week, as opposed to during the weekend. In red, are the number of monthly weekday users. In teal, are the number of monthly weekend users. Each month’s mean daily user count is mapped to the size of the circle and colored by day type. A local regression for each day type is shown to smooth the month-to-month effects and highlight the increasing trend.

A notable influx of MicrobeTrace usage occurred in late April 2020 (data not included in figure), simultaneously across nine cities in Vietnam over a span of two local afternoon hours. This brief influx of traffic from a single country, spread across disparate geography, is suggestive of workforce development efforts. If true, this would represent the first clear evidence of a training webinar held by non-CDC staff. Following on from this training event, the fraction of returning users was three times higher than MicrobeTrace’s historical fraction of returning users (64% versus 21%). Further, the average session duration was also nearly three times higher (20.1min versus 7.3min) than the historic average session duration.

## 4. Discussion

MicrobeTrace has been used to investigate a broad variety of infectious diseases. It has been used during CDC-assisted HIV cluster investigations in multiple states (Cranston, et al., 2019; Hogan, et al., 2017; John, et al., 2019; Shankar, et al., 2019), investigations of hepatitis C virus (HCV) (Falade-Nwulia, et al., 2018)), integrated into the Global Hepatitis Outbreak and Surveillance Technology (GHOST) that is used for viral hepatitis investigations (Longmire, et al., 2017) (S. Sims, personal communication), and is broadly used to integrate genomic and epidemiologic data for tuberculosis outbreak investigations (Springer, 2020). It has also been used to integrate partner services, epidemiologic and whole genome data to better understand transmission during a retrospective public health investigation of *Neisseria gonorrhoeae* (Town, et al., 2020). Outside of its intended domain of sexually transmitted diseases, MicrobeTrace has also been applied to integrate epidemiologic and laboratory data in outbreaks of foodborne pathogens, such as *Escherichia coli* O157:H7 (Allen, 2020). It is currently being evaluated for integration and visualization of epidemiologic and genetic data from cases of Ebola and COVID-19 (S. Whitmer, personal communication; S. Tong, personal communication).

MicrobeTrace offers a suite of capabilities to a public health expert that are typically only achievable with an array of software, tools, and custom scripts, and substantive computational experience. A putative MicrobeTrace user, such as epidemiologists or disease investigation specialist, typically achieves proficiency after one brief training session and aided by a cursory understanding of common browser interactions, such as ‘dropdown menus’, ‘slider bars’, and ‘drag-and-drop’. Many standalone tools are available to calculate pairwise genetic distances with varying degrees of specificity to the pathogen of interest. MEGA is a bioinformatic tool broadly used in public health, but new users can be overwhelmed by dense interfaces with scores of options that are often dense with jargon and required inputs (Kumar, et al., 2008). HIV-TRACE, which is specific to HIV sequence data, now offers rich visualization capabilities but its installation requires a keen understanding of Unix and the Git protocol for local installation and use (Pond, et al., 2018). An iteration of HIV-TRACE is available on the internet but at a web server which has concomitant data security issues (Weaver, et al., 2015). Patristic distance calculations are available via the APE package in R or the Java application PATRISTIC, but these require programming expertise and software installations (Fourment and Gibbs, 2006; Paradis, et al., 2004). Once genetic relationships have been calculated and contacts have been traced, integration and visualization of these links with individual-level data can be a complex task requiring tools like Gephi or Cytoscape (Bastian, et al., 2009; Smoot, et al., 2011). For those with programming expertise, integrated visualizations can be otherwise achieved with decade-old libraries in R with the iGraph package or in Python with the NetworkX and MatPlotLib packages (Csardi and Nepusz, 2006; Hagberg, et al., 2008; Hunter, 2007). Even so, these visualizations are not interactive with any additional figures, charts, tables, and maps that a public health expert might need to generate through the use of over a half dozen other applications (Figs. 2-4). If independently created, these visualizations must be augmented with network-level calculations and manipulations like threshold changes, minimum spanning tree calculations and filters, cluster membership, cluster size, and the number of neighbors for each node, all of which are easily performed in MicrobeTrace. These metrics can be manually calculated (e.g., R+iGraph, Python+NetworkX) or generated via opaque plug-ins in Gephi or Cytoscape that offer minimal customizations. Anecdotally, use of MicrobeTrace and its network layout interface can be playful; which has been shown to improve the user experience and increase their motivation to use the tool (Kuts, 2009).

While MicrobeTrace has been developed for a public health user base, it also has many applications in academia. It is adept at integrating arbitrary networks with independent node- and edge-level characteristics that are necessary to evaluate social, behavioral, biochemical, cellular, technological and physical networks. MicrobeTrace also offers rich customizations that reduce the time and effort to achieve insights and discoveries when grappling with a novel data set. The MicrobeTrace development team is not aware of another tool that offers all of these capabilities in a secure, interoperable, and light-weight format that requires no installation prior to use.

## Contributions

EMC and WMS contributed to design, project management, and manuscript writing. AB contributed to design, development, and manuscript writing. AS contributed to design, user manual, and manuscript editing. JK contributed to development. SK contributed to design and provided the nearest connected neighbor methodology.

## Acknowledgements

We are thankful to our colleagues in the Division of Tuberculosis Elimination (Kathryn Winglee, Sarah Talarico, Yuri Springer, Benjamin Silk), the Division of STD Prevention (Kim Gernert, Katy Town, Matthew Schmerer), the Division of Viral Hepatitis (Seth Sims, Garrett Atkinson, Yury Khudyakov), National Center for HIV/AIDS, Viral Hepatitis, STD and TB Prevention - Informatics Office (Max Mirabito, Silver Wang), Transmission and Molecular Epidemiology Team (Alexandra Oster, Cheryl Ocfemia, Nivedha Panneer, Scott Cope, Sheryl Lyss) for providing valuable feedback, features, bug reports, and continued training of our public health partners. We are also thankful to our user base in public health and academia for reporting bugs and suggesting features with regularity.

## Disclaimers

Use of trade names is for identification only and does not imply endorsement by the U.S. Centers for Disease Control and Prevention (CDC). The findings and conclusions in this report are those of the authors and do not necessarily represent the views of the CDC.

